# A lightweight segmentation and lineage tracking tool for noisy, low frame rate 4D fluorescence microscopy data

**DOI:** 10.1101/2020.11.01.364083

**Authors:** Haseeb Khalid Qureshi, Andor Magony, Yavor Hadzhiev, Kacper Wozniak, Alexandra Jasiulewicz, Ferenc Müller, Attila Sik

## Abstract

4D fluorescence microscopy allows the study of spatiotemporal cell dynamics in embryonic development in unprecedented detail, yet the uneven scattering of light within developing embryos presents challenges in discerning fine details. We present a tool which pre-processes large *in vivo* 4D microscopy datasets and can then track the movements and lineage histories of rapidly dividing embryonic stem cells. This solution offers a robust, simple segmentation technique to segment high intensity fluorescent features in highly scattered datasets, such as from deep within a developing zebrafish embryo. This tool then offers lineage tracing functionality by tracking rapidly dividing nuclei and their progeny, while also accounting for the jumps rapidly moving features make between frames in low frame rate data. Based on the user having prior knowledge of their imaging subject, this tool has applications in datasets with limitations with signal to noise, and frame rate. It can provide information regarding the position and movement of rapidly dividing nuclei. We demonstrate this pipeline in developing early zebrafish embryos.

## Introduction

High resolution laser light sheet microscopy has allowed us to observe whole embryological development in real time offering subcellular detail into embryogenesis. Fluorophores which bind to the nucleus are commonly used to visualise the embryo in development, offering a view into when and where cells divide. One such task which may arise from this is the tracing of origins and destinations of cells. Nuclei provide useful tracking anchor points as they are distinct and predictable cell organelles present in virtually every cell. Tracking cells (or their nuclei) can aid in the understanding of cell fates in early embryos, detecting heterogeneities in cell cycles and discerning global dynamics in cell division.

Tracking operates on converting imaging data into computationally trackable objects. To create objects out of foci of intense fluorescence, segmentation needs to be performed. It is here issues can be encountered, segmentation works best in cleanly imaged, high contrast, high signal to noise images (Dima et al. 2011). Such an ideal dataset is not always attainable.

### The imaging problem

Animal models such as the zebrafish, xenopus and drosophila are commonly used for studying embryonic development owing to the ease of access to embryos which are also highly suitable for microscopy. The early embryo undergoes a period of synchronised cell divisions, with progressive asynchrony over development. Zebrafish embryos offer a compelling model to study development *in vivo* due to the transparent nature of the embryos and their rapid development(Kimmel et al. 1995; Tadros and Lipshitz 2009).

A key drawback of *in vivo* imaging of a developing embryo is the scattering of light from deep within the sample. Fluorescence emitted from the centre of the embryo may be highly scattered, even under the most optimal conditions. The trade offs for deeply penetrative, high signal to noise imaging would remove the great asset of high temporal resolution which is offered by light sheet microscopy (in 3D imaging) as each frame would require more time to image. Such imaging may require either a greater laser power, or prolonged exposure time, or both. Prolonged exposure time will increase signal to noise but will compromise the temporal resolution, resulting in a longer time between frames, and potentially missing information in rapidly developing specimens(Keller et al. 2008). The prolonged exposure time to the laser light may also offer signal bleaching and phototoxicity (Jemielita et al.2013; Dima et al. 2011).

Low energy light sheet imaging is optimal for minimising frame times and phototoxicity. However, with some fluorophores this can increase the noise as long frame times and high energy produce cleaner images (Jemielita et al. 2013).

3D segmentation methods use slice by slice 2D segmentation, and then stitch 2D segments into 3D features. 2D segments can be created via the watershed algorithm. Temporal tracking of objects in 3D stacks provides 4D information of biological processes.

### The processing problem

The outputs of single channel light sheet imaging are a time series of 3D stacks, carrying 4 dimensions in the form of x,y,z,t (3D+t=4D). 4D datasets tend to be very large, ranging into the terabytes with regards to data size, so many current processing solutions are not optimal for general computing. Existing tools for parsing data of this nature include Fiji ImageJ, a highly versatile piece of software with excellent image file handling capability, but shortcomings in memory management due to its java based architecture (Schindelin et al. 2012), which may pose issues when trying to track nuclei in large datasets. Alternatively, tools like XPIWIT show great promise for tracking in early embryos, yet they involve complex pipeline building for data processing, and can struggle with high noise, low frame rate datasets (Bartschat et al. 2016). To create an accessible, scalable solution for tracking fluorescently illuminated nuclei in low frame rate and/or noisy images, we pursued a solution which will work on commonly available PC’s but could also utilise the horsepower of computing clusters.

### Introduction to our solution

The watershed algorithm is a commonly used segmentation tool, but popular iterations rely on basic thresholding to identify watershed seeds (Preim and Botha 2014). Given the uneven intensity topography throughout the embryo, we found that we needed an approach which did not rely on thresholding at all. This would enable it to adaptively identify candidate feature seeds based on local intensity topologies. Our solution used 1D intensity topography to identify intensity peaks from which to create seeds for the watershed.

This pipeline is designed for single channel grayscale images, ideally for nuclear stains although it can be appropriated for general blob detection and tracking (such as migrating cells in culture)

Our tracking solution takes the segmented nuclei and primarily seeks mitotic events to create lineages. Regional volume scanning via nearest neighbour analysis aims to detect the onset of mitosis based on the notion of mono nuclear cells.

Cell trackers commonly implement an overlap method where features need to overlap in space in consecutive frames to register as part of a track. Based on a method by Xiaowei et al 2006, our method compares Euclidean distances of nearest neighbours to determine the next point in a track (Xiaowei, Xiaobo, and Wong 2006), but is adapted to make use of knowledge regarding nucleus position in 3D space. This has use in low frame rate situations as spindle contraction during cytokinesis results in rapid movement of daughter nuclei, which will evade feature overlap methods of tracking as they move long distances between frames (Keller et al. 2008). A second feature detected in the scan space suggests a division has occurred, and measures have been taken to eliminate interference from false positives.

Embryonic stem cells undergo drastic volume decreases with successive divisions until a stability point where cell size cannot decrease any more (Reisser et al. 2018). This pipeline accounts for reductions in cell size, and potential stability points by offering parameter customisation by way of command line interface (CLI). Furthermore, our solution is lightweight so it can operate on general PC’s, owing to the step wise nature of how it approaches Z stacks.

Our segmentation/tracking pipeline shows promise in discerning characteristics of nuclei in dynamic settings, such as nucleus volume, topology and cell cycle length, and these can be tied to specific lineages, particularly in scenarios investigating cell fate. Furthermore, our method is optimised to work with single channel, single feature labelled images, removing the need for membrane dyes for discerning cell boundaries. This allows images using purely nuclear dyes to be tracked. However, as the core mechanic of nucleus detection relies on finding hyperintense regions of fluorescence, this is not restricted to nuclei. Such principles can extend to the tracking of other organelles, or even whole cells. With accessible source code, mostly built in Python, the tool is open to modification and modular design to suit individual needs, on top of the existing functions.

We demonstrate our segmentation and tracking pipeline in whole early stage zebrafish embryos, imaged using light sheet microscopy.

## Methods

The pipeline takes on a modular design with different functionalities compartmentalised into discrete, specialised scripts (**Fig 1**.). This allows customisation of pipeline use, allowing users to bypass unnecessary processing tasks for their specific use case. Sequential steps proceed from basic data acquisition to initial 2D binarization, 3D segmentation and to tracking. Quality control consists of inspecting outputs and modifying input parameters to optimise the pipeline, if data have been tracked incorrectly. For example, an insufficient scan radius may yield no mitotic events which is contrary to what a visual inspection of the raw data would reveal.

**Figure 1:**
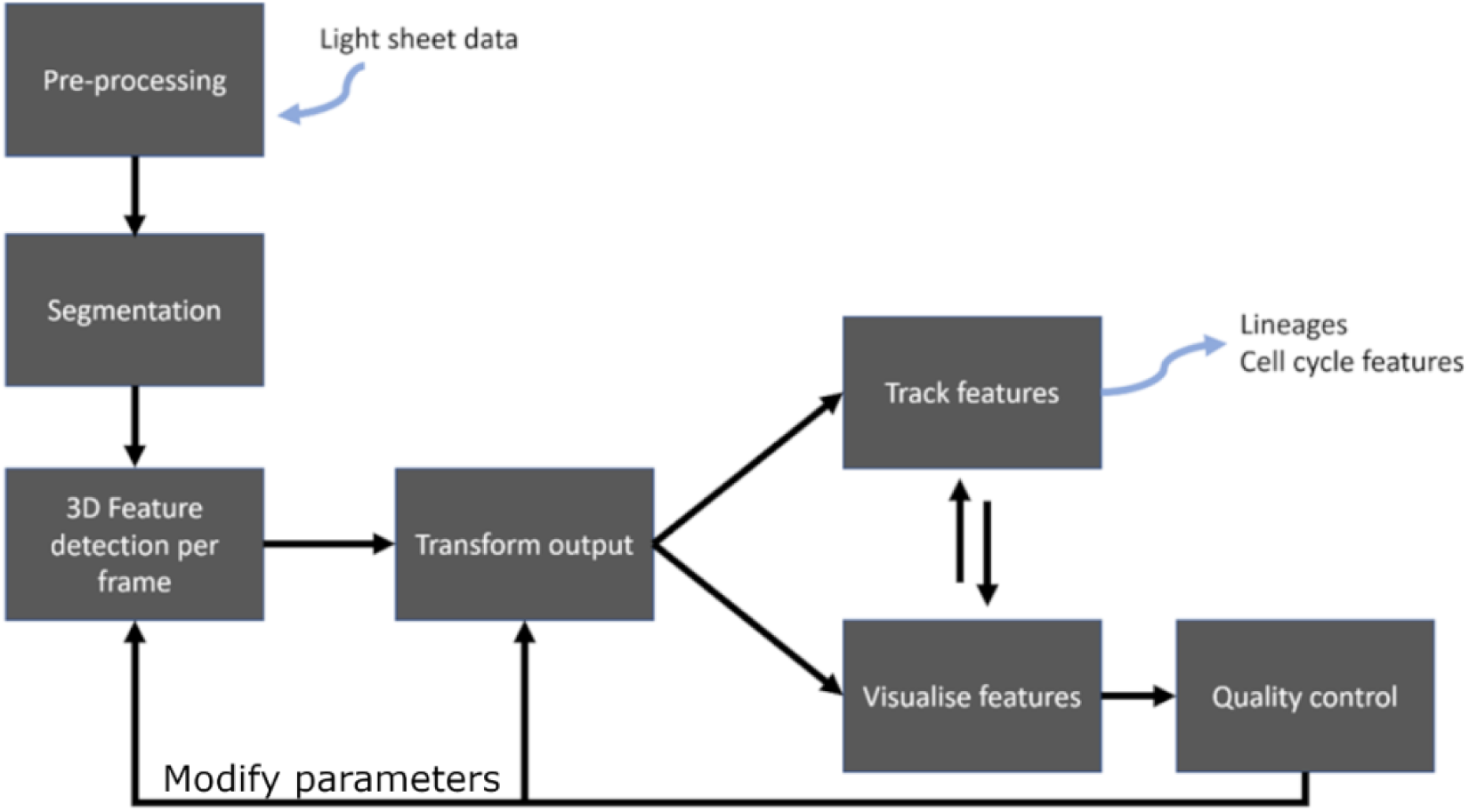
Architecture of tracking pipeline. Different modules are dedicated to different phases in processing.

### Imaging conditions

All data used to test the pipeline in this report were obtained using a Zeiss Z.1. laser light sheet microscope (Germany), using 20X W-Plan Apochromat water immersion lens with 0.91NA, 488nm laser (3% power). Samples were approx. 1-2 hours post fertilization (hpf) zebrafish (AB strain) embryos with nuclei visualised using Histone H2B MRuby injected into the yolk directly after fertilisation. Imaging took place at 28°C.

Segmentation operates on a Z slice by Z slice basis, thus works best with image files split into image series per time point. Each time point can be separated into individual folders, to parallelise processing of time points across processing threads and speed up processing time.

Single channel TIFF series work optimally, although other image file formats will work as long as one has the python libraries to process them, and one takes into account the bit depth of the image when setting peak detection parameters.

### Segmentation module

#### Thresholding based processing method

We explored standard image processing techniques for segmentation, and initially produced a MATLAB based (**Mathworks, MATLAB ver. 2019b)** pipeline based on the conventional watershed technique. For the sake of comparison, we will explore the steps taken to realise this method.

A median filter is applied to remove image grains and other undesirable noise patterns. At this stage, the image is cleaned of most artifacts and unwanted signals, ready for morphological transformations, and then converted to binary via thresholding

Small objects are removed based on a minimum size criterion for connected objects in the images. This size criterion is defined by the expected size of cell bodies, that has been obtained previously by manual image analysis.

This is followed by a labelling of 8-connected objects, and a Euclidean distance transform of the binary image. For each pixel in the binary image, the distance transform assigns a value that tells how far it is from the nearest nonzero pixel. The first watershed region identification is applied on the resulting image, after which we compute regional minima, that are connected pixel components with the same intensity value *t*, the external boundary pixels of which all have a greater value than *t*.

A morphological reconstruction is then used, so that the image only has regional minima where it is nonzero. This is followed by a repeated watershed algorithm. Contrast-limited Adaptive Histogram Equalization is then applied, which enhances the contrast of the image by transforming the values in the intensity image. It operates on tiles (small regions), rather than the entire image.

Image regions and holes are then filled by doing a flood-fill operation on background pixels of the binary image (imfill). Following this, the image is morphologically opened using an erosion function and dilation, with the same structuring element for both operations. Again, objects smaller than a specified number of pixels are removed.

#### Non-thresholding-based watershed seed generation (Fig 2a)

This is a Python based module which generates the binary representation for each Z slice in each stack. This module consists 2 key functions, which will be discussed in this subsection. The resultant binary masks are exported into the 3D segmentation module. This module requires user input on the following parameters before it can run:

**Figure 2:**
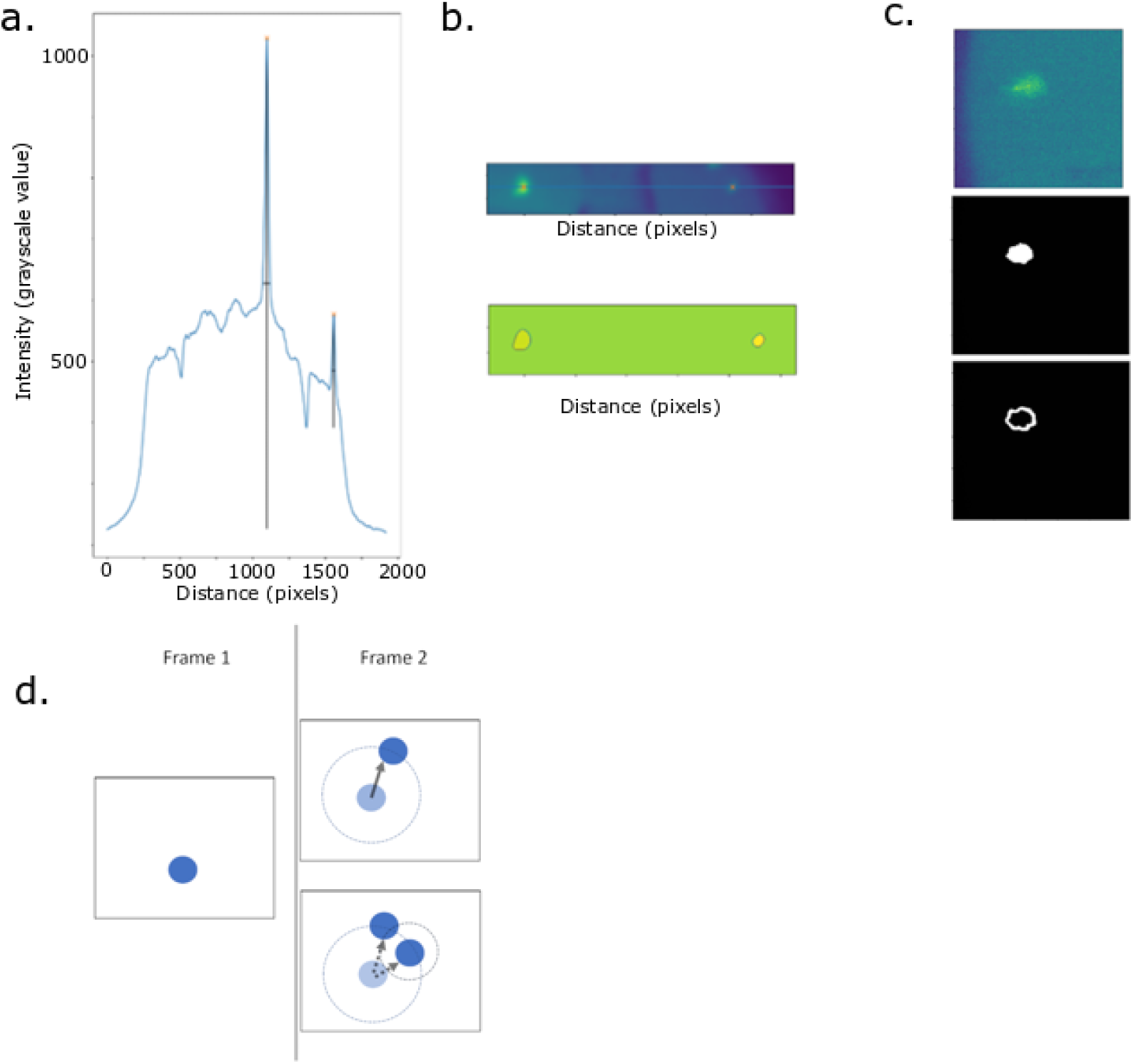
Illustrating key methodological steps in 2D segmentation and tracking. a: 1D feature detection by way of local intensity spikes. ii) Prominences are detected in 1D and iii) used to seed potential watersheds b: 1D feature detection hits from figure 2a, top) overlayed onto microscopy data. Bottom) resultant watershed using this seeding method. c: i) raw image, showing nucleus used for detection ii) watershed created on nucleus iii) watershed corona used to calculate foreground/background differential for retaining/discarding watershed d: Mitosis detection is achieved by comparing the location of a feature *a* at time point (frame) n (an), with the number of features within a defined radius of *a*_*n*_ in the frame n+1. If there is more than one feature, conditions are checked if this is a mitotic event. Frame 1 represents nucleus in current frame, frame 2 shows 2 scenarios. The top scenario shows when there is only 1 feature in the next frame within the scan radius (large dotted circle), indicating the continuation of the track. The bottom scenario shows a mitosis event, where 2 nuclei within a defined distance (smaller dotted circle) indicate the current track must end, and 2 new tracks must begin.

- x,y dimensions
- Maximum nucleus diameter
- Peak prominence
- Cluster distance
- Minimum 2D area of feature
- Maximum 2D area of feature

The watershed seeding layer is formed by detecting local intensity bright zones using the find_peaks function from the scipy library (Virtanen et al. 2020) (**Fig 2a**). The allows the tuning for peak prominence and allows for overdetection of watershed seeds from the selection of low peak prominence values. Peaks are detected on a 1D basis, as the algorithm scans the image matrix row by row. 1D scanning allows for granular peak detection, and 2D assessment of all detected peaks allows identification of clusters of bright zones which may constitute a feature (**2b**).

**User defined** peak prominence cut off intensity values (how tall a peak needs to be compared to its base), and which minimal intensity value to not consider peaks from. Indiscriminate peak selection can work better for detecting especially dim nuclei, but this will increase processing times

Detected peaks are represented in 2D space where DBSCAN clustering algorithm (from scikit-learn library) groups near peaks together(Pedregosa et al. 2011). **User input** is required to gauge the 2D pixel distance within which points are clustered together. Users are expected to input a maximal cluster distance, denoting the distance beyond which points are not linked to neighbours. These localised peak clusters represent potential features which will be discarded or retained based on the difference in intensity against the background.

These feature clusters are converted into watershed seeding points by scanning for the brightest peak in the brightest region of the cluster. Assuming the feature is a nucleus, the brightest region is expected to represent a more central point of the nucleus and provide an optimal seeding point.

#### Watershed refinement

The ‘basin’ into which the watershed seeds will spread is determined as a set circular zone around the watershed seed. The diameter of this is user defined as the expected maximum nucleus diameter. In early developing embryos, the nucleus and cellular diameter are expected to decrease dramatically, so a maximal diameter is considered to be all encompassing for future cell diameters.

With the watershed seed and unknown zone generated, watersheds can now be created. Watersheds of nuclei are generated using the Python OpenCV2 library. Once watersheds are competed, each is processed by dilating the watershed by a factor of 10% and comparing the average intensity of the corona produced by dilation with the average intensity of the watershed (**Fig 2c**). If the differential is less than the default 10% (this can be adjusted to match specific data dimensions) then it is discarded. These watersheds are saved as a binary representation to feed into the 3D segmentation portion of the pipeline.

#### Parallel Processing

The segmentation is parallelised across a user defined number of logical CPU cores. Each time-point corresponding folder is processed sequentially, but the contents of each folder (the z slices) are distributed across cores and queued for parallel segmentation.

#### 3D Segmentation module

This module is a MATLAB based 3D segmentation tool which takes the binarized Z slices (per time point) and constructs 3D features out of them. This outputs 3D geometrical data by way of Excel csv files for each time point. These files contain information on the coordinates of each feature centroid, as well as x,y,z dimensions and volume. This step measures the properties of image regions with shape measurements (e.g. ‘BoundingBox’, ‘Centroid’, ‘Area’, regionprops), after which a database can be built containing detected cell parameters for further analysis and for cell tracking purposes.

Though offering caveats in memory management for full z stack processing, MATLAB proves effective in 3D visualisation and segmentation capability. The 3D visualisation function can be used on a single sample stack to test the outputs are processed correctly, then can be switched off to allow for more memory efficient 3D segmentation.

#### Transformation module

This is a Python based script to bridge the 3D segmentation output with the tracking output. The centroids of the 3D features created via the connected components method are stored in a csv file, alongside their volumes and x,y,z dimensions. A transformation script converts x,y,z dimensions to microns, via user input stating the conversion factors, and these are read by the tracking script.

#### Tracking module

This is a Python based module which takes the centroid data and joins them into tracks and lineages based on spatiotemporal proximity. The tracking module requires user input on the following parameters before it can run:

- Data length (frames)
- Maximum expected scan radius
- scan radius decay factor (an estimate of how much the cell radius is expected to decrease per cell cycle)
- mitosis buffer (number of frames after mitosis when division is not expected)
- minimum expected scan radius

The tracking script begins tracking from the first time point (time point 0), and stores all centroids from time point 0 as starting points for tracks. Each point is taken and is compared with all centroids in the next time frame to find the nearest centroid. This is achieved by using nearest neighbour analysis from scikit-learn library. The nearest neighbour analysis only scans for centroids within a **user defined radius**. This radius is expected to represent a volume less than the radius of the entire cell, as cytokinesis sees rapid migration of daughter nuclei to opposite poles of the cell.

This section of the tracking takes into account loss of cell and nuclear diameter volume with successive divisions in early embryos so has a **user defined decay factor**, and a plateau point beyond which one does not expect cell diameter to decrease notably anymore. This is an important parameter as too large of a scan radius may continually trigger mitosis events and perpetually run the program.

If the nearest neighbour function detects multiple centroids within the scan radius in the next frame, this may suggest that mitosis has occurred. The algorithm searches for the nearest feature within the scan radius, and then the nearest feature to that within a smaller scan radius defined by a smaller user defined by a smaller user defined radius (also influenced by the afore mentioned decay factor) (**Fig 2d**). If a sister nucleus cannot be located within this radius, the next nearest figure is checked for these same parameters. If this also fails to yield a sister nucleus, the track is halted to eliminate tracking of false positives, and to keep lineages as whole as possible. If two suitable features are detected, the current track is terminated, and 2 new tracks are created. These tracks are added to the time point 0 origin pool, and will be tracked as usual, except they carry the metadata of their lineage. This metadata records how many divisions have occurred before, and which half of the mitotic daughter pair they are.

This is continued until there are no points left in the tracking origin pool. The output produced is a matplotlib 3D line plot showing a 3D representation of tracks, colour coded to show different generations.

#### Validation

Validation of tracking parameters was achieved by manually identifying mitotic events in a maximum intensity projection of the raw data, and comparing with the timings of mitotic events from the tracking tool. Numbers of nuclei per frame were counted, and the rise in detected nuclei was also measured. This validation allows us to manually compare cell cycle lengths with the tracking tool, as well as measure when the rise in detected features occurs (suggesting the multiple occurrences of mitosis within the specimen).

Additionally, a temporal maximum intensity projection was created (max projection in T, rather than Z) using Xen Black and was input into a script where the mayavi library was used to overlay the tracked paths of detected nuclei with a static 4D representation of the tracked image. This allows the user to visualise how the tracks line up with the raw data.

## Results

### The intensity topography of the early embryo

We initially sought to understand how fluorescence intensity varies around the embryo, in order to find out why consistent segmentation was so difficult.

Using conventional watershed seeding methods from MATLAB revealed issues with the uneven intensity topologies in noisy light sheet data (**Fig 3b top**). Taking 1D histograms across z slices at various depths in an embryo shows light scattering in the embryo leads to high signal to noise imaging on the edges of the embryo but a more uniformly bright interior (**Fig 3a**) presenting a challenge to image processing. As embryonic development progresses and fluorophores evenly distribute or become bleached, the intensity topography shifts. This led to problems with thresholding (even after application of histogram equalisation and median filter to even out noise and intensity) the embryo for watershed, and stimulated the development of a more robust watershed method which would produce binary images to feed into 3D segmentation for tracking.

**Figure 3:**
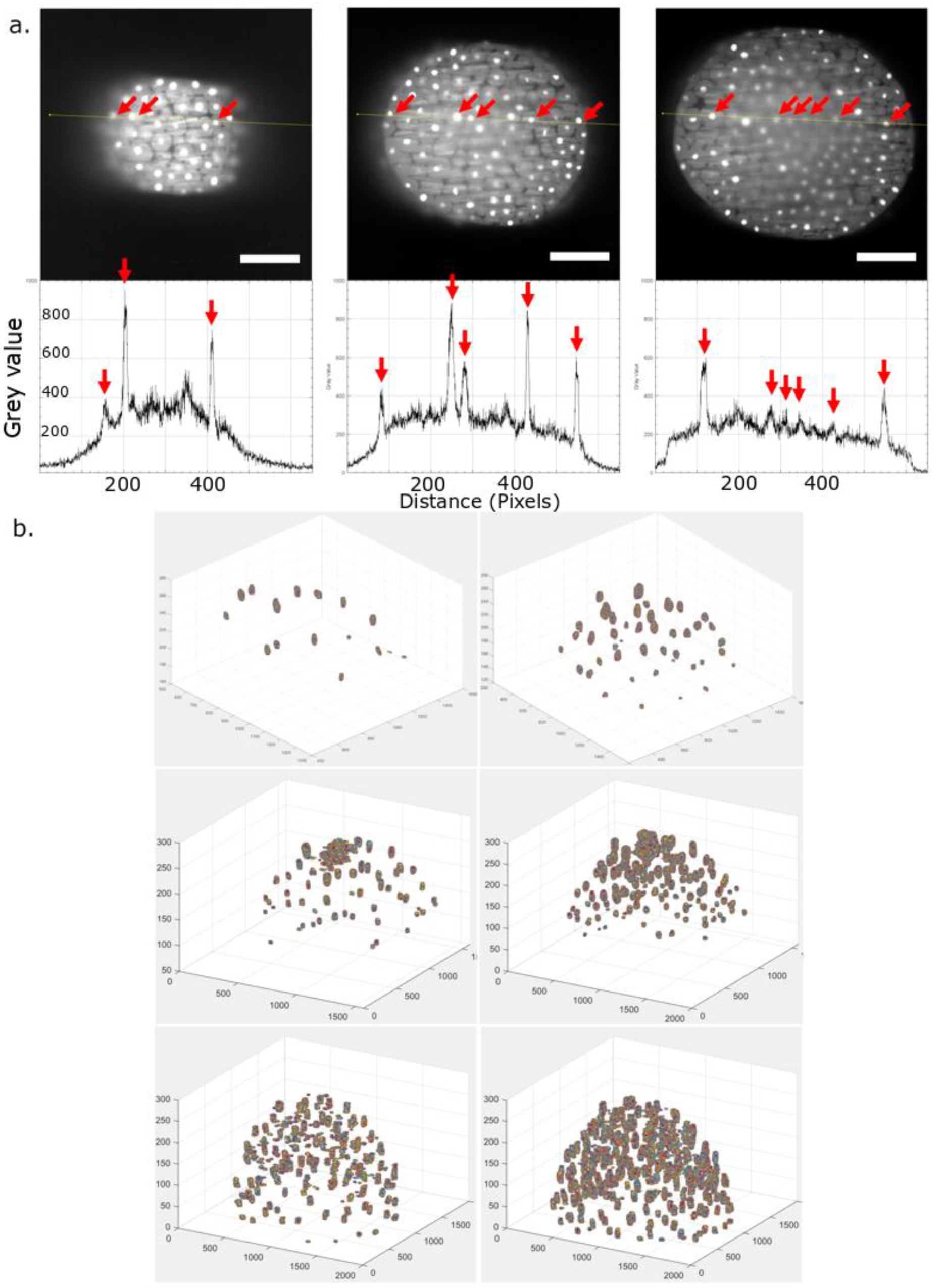
Navigating the dynamic intensity topography of a developing embryo. a: 1D intensity topography (taken along yellow line) of different z slices (upper panels) from progressively deeper (left to right) regions of embryo. Red arrows denote nuclei, and are lined up between the z slice and their graphs. b: Consequences of 3D segmenting binary outputs of differing watershed methodologies. Left panel is segmentation of data from 128 cell stage and the right panel is at 256 cell stage (same embryo, same time lapse) **top)** underthresholding can lose many features whereas **middle)** Overthresholding to capture more features can capture swathes of brightly illuminated zones as a single feature **bottom**) shows the 3D segmentation outputs applied on the binary outputs from our novel segmentation technique

We noted the uneven intensity topography of the embryo presented challenges to segmentation involving a global threshold (**Fig 3b**). Using a more conservative thresholding value would underthreshold the image. The 3D segmentation data shown in **Fig 3b (top left panel**) showed 16 nuclei at 128 cell stage of the embryo, and the same setting applied later in the time lapse yielded 61 features at 256 cell stage (**Fig 3b top right panel**). Lowering the threshold to raise inclusivity of features presented an overthresholding problem, whilst still underreporting the number of nuclei yielding 47 features at 128 cell stage (**Fig 3b mid left**)and 163 features at 256 cell stage. While the number of features detected has increased, there is no reason to continue decreasing threshold as multiple features are merging into a single one, as the intensity at the top of the animal cap is generally greater than the intensity in the middle of the embryo (**Fig 3a**). Running the binary outputs from our novel 2D segmentation method (**Fig 3b bottom**) yields 154 features at 128 cell stage (**left panel**) and 292 at 256 cell stage (**right panel**). This indicated detection of noise, which may arise from fragmentation of nucleus during segmentation, or extreme fluctuations in local intensity topography being accepted as a nucleus by the algorithm.

These data aim to show the dynamic intensity environment within specimens, and the obstacle conventional image processing methods must overcome.

### 2D segmentation algorithm

With the understanding of how the intensities of features and their local environments varied in imaged specimens, we sought to design an alternative segmentation technique which relied on local prominences and relative contrast against background noise.

Implementing the steps in the segmentation step reveals nuclei in the embryo as hyper intense foci which show a steep difference in intensity compared to their immediate surrounding (**Fig 2a, Fig 4a, 4b**). This concept is utilised in watershed refinement. As is clear in **Fig 4c**, peak detection detects many intensity peaks dependent on the signal to noise of the image. Watershed refinement removes erroneous features (**Fig 4h)** although quantification of the feature detection and watershed retention rate reflects greater retention of features with increased depth. Manually counting the retention of detected features versus the actual number of features suggested a detection rate of between 85% and 99% in the deep, highly scattered embryo (**as represented in Fig 4g right**). The fidelity of the watershed refinement function relies on setting an appropriate corona to feature differential, which can be determined by testing the script on a small sample of slices representing the noisiest region of the specimen and iterating until maximum nuclei are detected.

**Figure 4:**
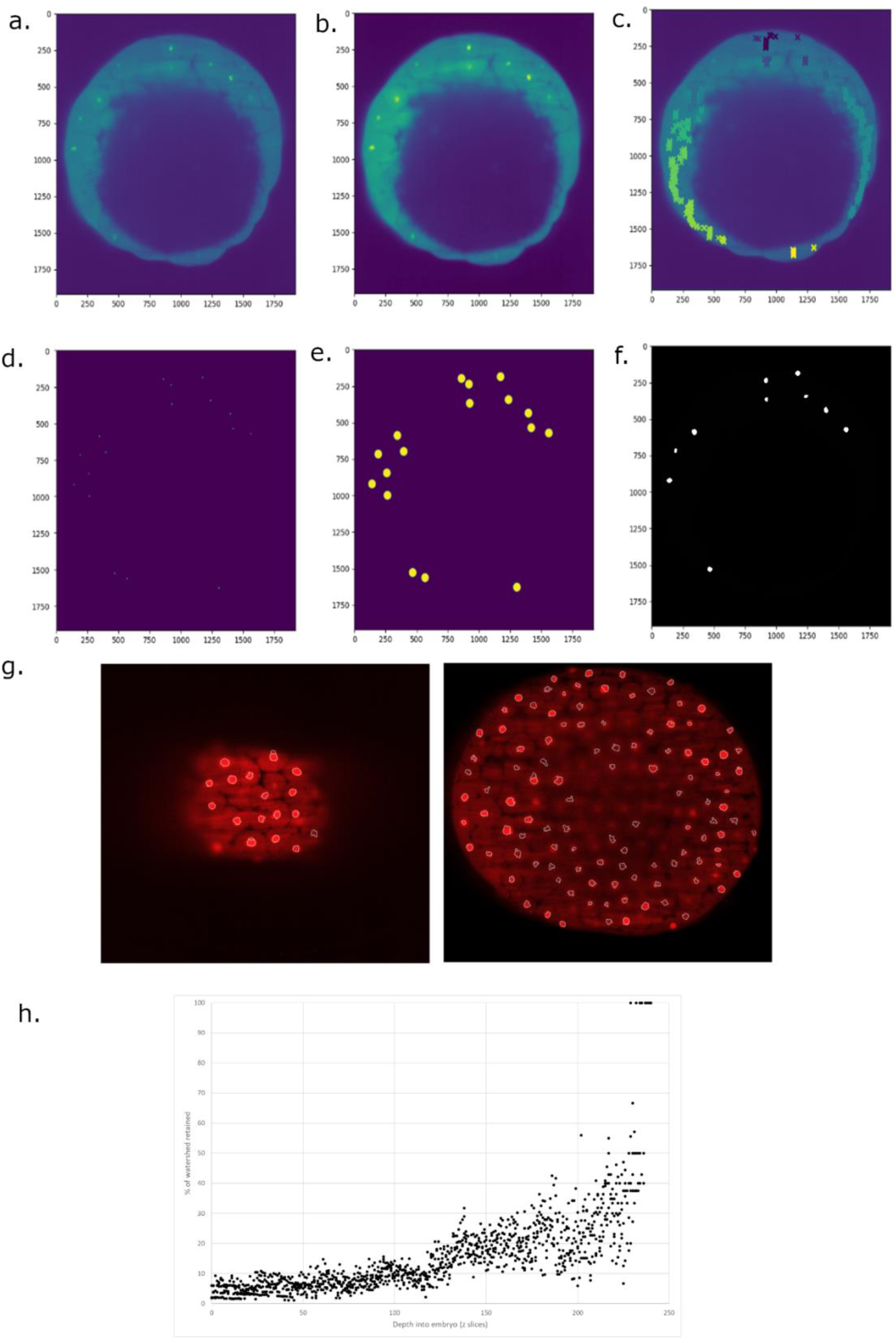
2D segmentation methodology and visualisation. **a-f** show segmentation steps in zebrafish embryo at region where yolk enters frame (large ‘hole’ in image). Axes show size of image in pixels. x,y Pixel/micron conversion is 1px = 0.33 microns **a**) raw image **b**) gaussian blur **c**) groups of watershed seeds detected by peak detection and clustering **d**) refinement of peaks to most central bright point **e**) background elimination for watershed seeds to scan into **f**) refined watersheds (after removal of false candidates) **g**) Nucleus detection (white outlines) in clear surface region (left) and scattered deep region of embryo (right) **h**) Feature retention rate from the watershed elimination function, showing proportion of features kept by watershed elimination per descending z slice: likely nucleus candidates. Data from multiple frames in same embryo.

Notably, detection of nuclei in noisy regions may miss the polar tips of the nuclei, yet 2D segmentation will likely still capture the bulk of the nucleus in the interceding z slices, allowing a centroid for that nucleus to be created, maintaining use for tracking.

We have shown a breakdown in the steps taken to segment hyper intense foci (in this case corresponding to nuclei), and how this can function in regions of highly scattered light.

### Tracking algorithm

Using the built-in function to identify mitosis events and generate family trees from tracks, we are able to produce clear family trees of nuclei, which can be tailored to represent relevant metadata. In this case, the length of each branch corresponds to cell cycle length. The outputs from tracking are in accessible csv files, so are open for bespoke analysis depending on a users needs.

Lineage tracing appears to follow expectations, following trends in embryonic mitotic waves, and closely matching the expected number of features (**Fig 5a**). Cell cycle lengths matched manually tracked cell cycle lengths (anaphase to anaphase), with no significant differences between manually tracked nuclei and automatic ones, once correct tracking parameters had been chosen and noise discarded (**Fig 5b**).

**Figure 5:**
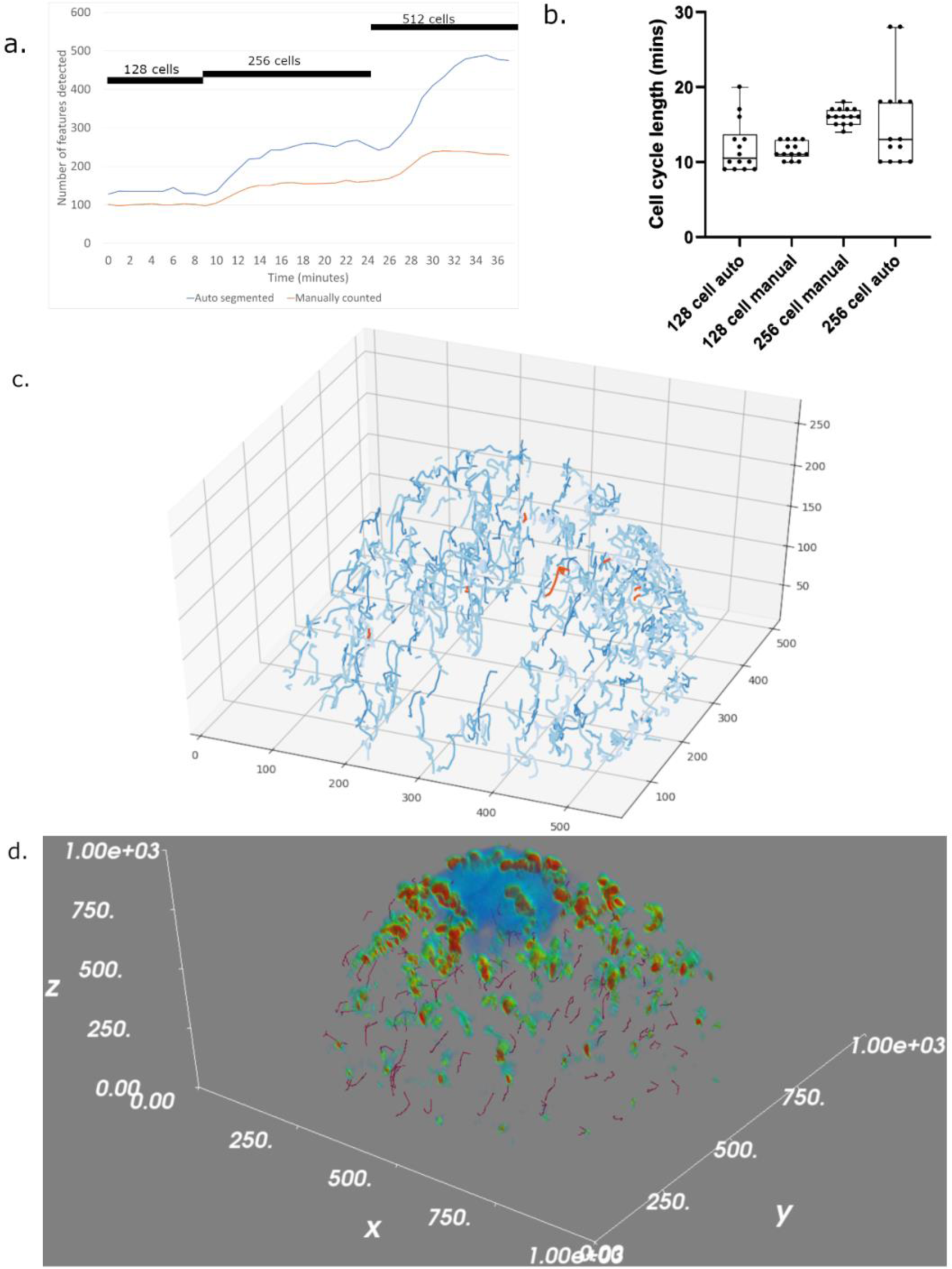
Outputs from tracking nuclei in a developing embryo, and tracking validation. a: Comparison of detected features between manual counting (using 2D max projection) and 3D segmentation via our pipeline, showing the increase in features with synchronised waves of mitosis (denoted by black bars). b: comparison of tracked cell cycle lengths between manually tracked and tracked automatically via the pipeline. c: Tracking outputs shows the lineages tree in 3 Dimensional space, with lighter hues representing successive generations. d: Mayavi based validation tool, overlaying the tracks with a 4D maximum intensity projection. Only the first cell division is represented in the image rather than the full set of lineage tracks.

In high frame rate data sets, the scan radius need not be so large as there is less chance of missing the daughter nucleus in the next frame, but in low frame rate data sets the daughter nucleus may have moved across the cell before the capture of the next frame. In these cases, a larger scan radius is required. It is helpful to take a glance at how far ones own nuclei move following a cell division, and using that base this information on. Optimising parameters (the quality control step in Fig 1) will produce a network of lineage trees plotted in 3D space (**Fig 5c**). This data can be extracted from the resultant csv files and used for lineage analysis.

In a dataset where expected cell cycle lengths were known, we noticed overtracking begins when there is a sudden increase in abnormally short cell cycle tracks. We noted that another key indicator that overtracking is occurring is a drastic jump in tracking time, or even an indefinite tracking loop.

It is wisest to test the program with a smaller scanning radius: using nucleus diameter + 10% and increase the diameter (and adjust decay factor) until the detection output begins producing many incorrectly short tracks. Such short tracks would be far shorter than a cell cycle possibly can be, this suggests tracks are ending prematurely due to too large scan radius triggering mitosis events, or it may just be a high number of false positives interfering with tracks.

Overlaying tracks with a 4D representation of the timelapse allows direct validation by viewing the tracks in relation to the actual data (**Fig 5d**). This step can allow one to identify of the tracking parameters selected need tweaking. Manually counting nuclei in a maximum intensity (in Z) projection timelapse shows the detection rates of features aligns with the expected number of nuclei to detect. Cell cycle lengths from the tracking algorithm show agreement with the manual tracks, and the timings of the synchronised mitosis events also appear to align with the manual measurements. It is notable, the tool picks up additional noise and tracks that too. We would recommend filtering for size in the transformation tool, or in the case that tracks seem to run for very long, increasing the scan radius may also improve track quality (at the expense of processing time).

Through these tests, we demonstrated our tracking algorithm, and how the outputs can be parsed and validated.

## Discussion

The capabilities of our tool reflect the specific issues our laboratory investigates, yet this tool is designed with general use in mind. The ethos behind the design of this workflow was to perform single tasks as effectively as possible. The various modules can work as stand-alone programming modules to perform their individual tasks, although their inception necessitated them to be used in a pipeline designed to segment and track nuclei in a rapidly dividing *in vivo* model.

The segmentation module allows for consistent nucleus detection across the 3D of the intensity topography of the embryo. It is able to discern nuclei from the highly scattered optical environment of the deep embryo, with quantitation suggesting more favourable segmentation in the highly scattered environment deeper into the embryo. We still receive favourable detection rates in brightly illuminated surface regions, yet the sensitive feature detection algorithm simply offers more noise for the watershed refinement tool to remove. This may impact processing times.

The lineage tracing tool offers granular information on the changes in cell size and cell cycle length, making the tracking tool a useful base for identifying lineage specific characteristics.

Analytical capabilities regarding spatiotemporal features of mitosis events (specifically cytokinesis), allows quantification of cell division coordination. In particular, there is the scope to investigate coordinated ‘waves’ of mitosis in early embryos (Anderson et al. 2017; Vergassola, Deneke, and Di Talia 2018).

The wider use of this pipeline translates to investigations which involve isolating lineages to high noise 3D datasets. The threshold free segmentation algorithm is robust, and has utility as a binarization step in other image processing scenarios. The standalone nature of the segmentation module gives it portability to use in applications which may not involve lineage tracing.

This pipeline, though designed to work around shortcomings in acquisition of microscopy data, is still limited by raw data quality. Whereas a key strength of this tool is its ability to register the presence of a nucleus, the precision with which the watershed discerns feature geometries is still influenced by light scattering. We found the core of the nucleus is reliable detected, but the edges may not always be detected correctly. This drawback makes volume measurements only as good as the image quality, but the integrity of the tracking appears to be mainly intact. Maximising accurate watershed may rely on pre-processing, or in the worst case, changing imaging protocols. We also find that the use of a scan radius optimises this for use in specimens where cell size is broadly similar. Despite being tested in 3D embryonic datasets, these principles can translate for use in cell culture. The algorithm will simply process it as if it’s a 3D stack of 1.

By comparing with more conventional image processing methods, our novel 2D segmentation method uses fewer image transforms on the raw data, and focuses more on the user tailoring the parameters of the segmentation algorithm to match the dynamically changing fluorescence environment of the specimen. This provides accessible, granular control of a computer vision framework to pick out points of interest in 3D microscopy data.

Finding the optimal parameters requires some experimentation, so we would recommend taking an early time point and a late time point and testing the segmentation parameters on these data before running the program on the entire dataset. The parameters which have the biggest impact on segmentation appeared to be the corona differential, and the peak prominence parameters. Other parameters simply rely on basic knowledge of one’s own dataset, such as a rough estimation of the size of a nucleus at the beginning and end of the dataset.

It helps to have an understanding of the biological system, such as the average cell cycle length (as you can identify if your tracking parameters need refining), but a risk with this is that one may just influence the data to meet their expectations and potentially lose interesting novelties. As a result, it is possible to navigate the outputs of this software without adhering to prior knowledge of expected cell cycle lengths.

## Conclusion

This pipeline is suited to developmental biology problems where big data can be prohibitive to analyse. This offers a computationally lightweight, streamlined approach to segmenting and tracking nuclei in rapidly dividing embryos (either low frame rate temporal data where there is no geometric overlap of features between frames, or simply quickly dividing nuclei). The approach to watershed seeding also makes the segmentation a powerful standalone tool for segmenting in noisy data, although precision of feature geometry is still somewhat dependent on signal/noise ratio.

## Acknowledgments

This work was supported by the European Union’s Horizon 2020 Marie Sklodowska Curie Research and Innovation Staff Exchange programme (VISGEN, No. 734862) and OPEN FET RIA (NEURAM, No, 712821), the Higher Education Institutional Excellence Programme of the Ministry for Innovation and Technology in Hungary, within the framework of the “Innovation for the sustainable life and environment” thematic programme of the University of Pecs, and the Wellcome Trust Investigator award to FM.

